# What controls carbon sequestration in plants under which conditions?

**DOI:** 10.1101/2023.04.20.537672

**Authors:** Tim Nies, Marvin van Aalst, Nima Saadat, Josha Ebeling, Oliver Ebenhöh

## Abstract

Plants use photosynthesis to harvest sunlight and convert the solar energy into chemical energy, which is then used to reduce atmospheric carbon dioxide into organic molecules. This process forms the basis of all life on Earth, and stands at the beginning of the food chain which feeds the world population. Not surprisingly, many research efforts are currently ongoing aiming at improving plant growth and crop yield, and several of these activities directly target the photosynthetic pathways. Metabolic Control Analysis (MCA) shows that, in general, the control over a metabolic flux, such as carbon fixation, is distributed among several steps and highly dependent on the external conditions. Therefore, the concept of a single ‘rate-limiting’ step is hardly ever applicable, and as a consequence, any strategy relying on improving a single molecular process in a complex metabolic system is bound to fail to yield the expected results. In photosynthesis, reports on which processes exert the highest control over carbon fixation are contradictory. This refers to both, the photosynthetic ‘light’ reactions harvesting photons, and the ‘dark’ reactions of the CalvinBenson-Bassham Cycle (CBB cycle). Here, we employ a recently developed mathematical model, which describes photosynthesis as an interacting supply-demand system, to systematically study how external conditions affect the control over carbon fixation fluxes.

## Introduction

Photosynthesis has been classically divided into two parts. The ‘light’ reactions supply energy and reduction equivalents to the ‘dark’ reacions of the CalvinBenson-Bassham (CBB) cycle [1, 2], where carbon dioxide is fixed to form reduced carbon compounds used as building blocks in other metabolic processes. The CBB cycle (demand side) is one of the most critical pathways on earth that plants and many other photosynthetic organisms use. Current estimates in-dicate that over 99% of global carbon dioxide is fixed by the key enzyme of the CBB cycle, ribulose-1,5bisphosphate carboxylase/oxygenase (RuBisCO) [24]. To guarantee efficiency and prevent the formation of toxic reactive oxygen species, the supply and demand of energy and redox equivalents must be coordinated [17]. However, the habitats of photosynthetic organisms are usually characterized by a high fluctuation of abiotic factors, such as light intensity and CO_2_ concentration [16], which makes balancing the photosynthetic electron transport chain (PETC, supply side) and the CBB cycle challenging. Therefore, versatile regulatory mechanisms that coordinate carbon fixation and the PETC and adapt both processes to external conditions have evolved. Examples of regulatory mechanisms include non-photochemical quenching, the thioredoxin dependent redox control of CBB cycle enzymes, and regulated changes in stomatal conductance [6, 19, 7]. All of these processes are currently targets of research activities aiming to increase plant performance and crop yield [15].

Considering the importance of the PETC and CBB cycle, it is unsurprising that much effort has been spent studying their kinetics, regulation, and control by experimental and theoretical methods. Various mathematical models have been developed aiming at providing a theoretical framework to analyze which factors determine the efficiency of carbon fixation [8, 9, 20, 21, 12]. Kinetic models of the CBB cycle established, e.g., the importance of the sedoheptulose1,7-bisphosphatase (SBPase) for controlling carbon assimilation and provided theoretical explanations for a wide range of observed kinetic properties of RuBisCO [21, 23, 33]. Appropriate theoretical tools are needed to study control in metabolic networks, e.g., the CBB cycle and PETC. Metabolic control analysis (MCA) is a theoretical framework developed in the 1970s, which is continuously improved and generalized [10, 14, 11, 4, 32]. A major purpose of MCA is to quantify the influence that single enzymes have over steady state properties of metabolic networks. A central concept are the *control coefficients* which describe how small changes in activities of single steps affect stationary metabolites and fluxes. Because control coefficients depend on the dynamics of the interactions of all components, they are systemic properties. MCA has been repeatedly applied to study the control of reaction steps in plant metabolic path-ways [25] with examples including applications to the benzoid pathway, sucrose accumulation, the CBB cycle, the electron transport chain, and combinations of these [30, 3, 5, 17, 26].

Here we present an *in silico* analysis of how external conditions affect the control over carbon fixation fluxes. We focussed in particular on the effect of two environmental parameters, light intensity and CO_2_ concentration, to assess how these factors affect the control of carbon fixation. Classically many studies stress the importance of RuBisCO and its activation processes as highly influential on photosynthetic efficiency [28]. But is RuBisCO always the main controling factor? For our analysis, we employ a published kinetic model of photosynthesis that combines the PETC, the CBB cycle, and the AscorbateGluthatione (ASC-GSH) cycle [26]. This model was originally used to study the importance of cyclic electron transport around photosystem I for photoprotection, and also includes regulatory mechanisms, such as non-photochemical quenching, state transitions between the two photosystems, and redox regulation of CBB cycle enyzmes through the thioredoxin system. We began our analysis by using the stoichiometric structure of this combined model to conduct a reaction correlation analysis [22], followed by the investigation of control on carbon fixation by different key processes in photosynthesis. We identified a condition-dependent shift in control and determined its structural origin using a robustness analysis with sampled parameter sets. Using the results from the robustness analysis, we could show that some reactions exert control in an either-or relationship while others may exert control simultaneously. With this work, we contribute to elucidate the control in photosynthesis and its dependence on external conditions.

## Results

### Reaction correlation analysis

We begin our analysis by studying the constraints on stationary fluxes, which are imposed by the stoichiometry of the network alone. For this, we calculate reaction correlation coefficients [22] for the previously published photosynthesis supply-demand model [26]. These coefficients provide a generalization of the concept of enzyme subsets. Reactions within one enzyme subset are strictly coupled in the sense that they always carry fluxes in a fixed proportion. For such reactions, the reaction correlation coefficient is *±*1, and the corresponding row vectors of the kernel matrix of the stoichiometry matrix are parallel. Reaction correlation coefficients generalize this idea by essentially calculating the angle between row vectors of the kernel matrix (for details, see [22]), and therefore indicate how strong reactions are correlated as a result of structural constraints of a network. Fig. 1 presents a metabolic tree constructed by hierarchical clustering using dissimilarities calculated by the reaction correlation coefficients (see Methods). The matrix of reaction correlation coefficients *ϕ* can be found in the supplement (Fig. S1).

**Figure 1.**
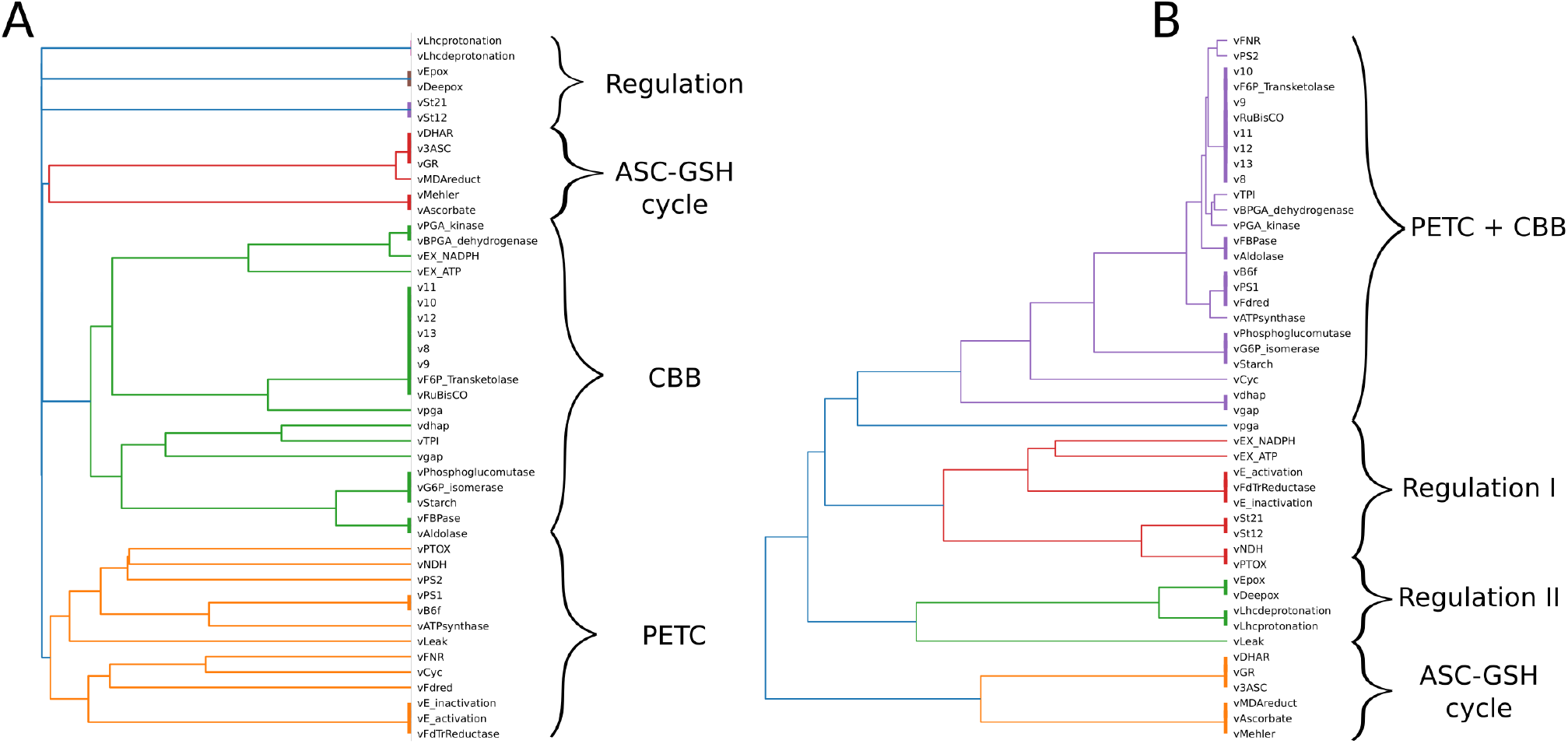
Metabolic trees, showing different metabolic clusters. The left tree was obtained by reaction correlation analysis. The right tree was obtained by steady state flux correlation analysis for 10000 random parameter sets in low light intensity and low CO_2_ concentration conditions for the kinetic model of photosynthesis. Abbreviations in the cluster annotations: PETC — photosynthetic electron transport chain, CBB — Calvin Benson Bashham cycle, ASC-GSH cycle — ascorbateglutathione cycle.

There exist ten enzyme subsets containing more than one reaction. Generally, reactions that function as regulatory control mechanisms (such as deprotonation / protonation of PsbS, and the xanthophyll cycle necessary for non-photochemical quenching, or the reactions involved in the redox control by thioredoxin), each form an enzyme subset. Three subsets are connected to the electron transport chain and ROS scavenging. The first subset consists of the reactions mediated by cytochrome *b*_6_*f* and photosystem I, which are therefore strictly coupled. The second contains the Mehler reaction and ascorbate peroxidase, while the third consists of the remaining reactions of the ASC-GSH cycle. Four subsets can be assigned to the CBB cycle and starch synthesis. The largest subset contains eight reactions, including RuBisCO, the transketolase reactions, and the reactions of the regeneration phase leading to the formation of ribulose1,5-bisphosphate.

In Fig. 1, clusters of reactions, which are highly correlated but not strictly coupled, are indicated by different colors. We identify clusters that are associated with key metabolic functions, such as the PETC, the ASC-GSH cycle, and the CBB cycle. The CBB cycle is split into two pronounced subclusters, which can be associated with carbon fixation and carbon export, respectively. The same is true for the PETC, for which we can distinguish between the linear (photosystems II/I and cytochrome b_6_f) and the cyclic electron flow. Interestingly, RuBisCO and PSII are grouped in different clusters, which suggests that these two processes are decoupled. This is unexpected because the electrons obtained by photosystem II are mainly used by the CBB cycle to fix carbon, therefore, one would expect highly correlated fluxes. This observation can be explained by considering that the stoichiometric analysis applied here is based on the full nullspace, and completely ignores constraints on the stationary flux solutions by the kinetic parameters

A kinetic model, however, drastically restricts the possible stationary fluxes by the specific parameter values of the dynamic equations. Therefore, actually observable steady-state fluxes will represent only a small subset of the complete nullspace. The processes decoupling reaction rates of RuBisCO and PSII, such as proton leak, terminal oxidases, etc., are constrained by their kinetic parameters to carry only relatively small fluxes, such that, in fact, the correlation between RuBisCO and PSII rates should be highly correlated for realistic conditions. To test this assumption, we repeated the correlation analysis based on steady-state fluxes sampled from the kinetic model, in which the reference values of the rate constants have been randomly varied by a factor between 0.5 and 2. The rationale behind this is that now the stationary fluxes are restricted to solutions which are close to a reference state and therefore reflect a more physiologically relevant subset of the nullspace. The resulting tree is depicted in Fig. 1B. As expected, the fluxes of the CBB cycle and the PETC are now strongly correlated.

### The control on carbon fixation switches between environmental conditions

We use metabolic control analysis (MCA) to quantify the control that individual molecular processes exert on the performance of the system, measured by the net carbon fixation rate, in different environmental conditions. We found that overall flux control is exerted mostly by one of four steps: photosystem I and II in the electron transport chain and RuBisCO and SBPase in the CBB cycle. In Fig. 2 we depict the four flux control coefficients on the overall carbon fixation rate for light intensities ranging from 50 to 1000 μmol m^−2^ s^−1^and for CO_2_ concentrations between 6 and 20 μM, corresponding to atmospheric concentrations between approximately 170 and 700 ppm. It is clearly visible that there are two distinct light regimes. A sharp transition between control by the light reactions (low light) to control by the dark reactions (high light) can be observed. Interestingly, the curve separating these two regimes corresponds to the limit where the quenching capacity reaches its maximum, and the lumen becomes highly acidic (see Fig. S2). As we observed and discussed previously [26], this transition marks the saturation of the photosynthetic system, above which increasing light no longer facilitates higher carbon fixation rates. At the transition, carbon fixation rate (and many other rates and intermediate concentrations) is not a smooth function of the incident light intensity. Therefore, the numerical differentiation employed to calculate the control coefficients may lead to imprecise results and as a consequence the coefficients very close to the transition should be interpreted with care.

**Figure 2.**
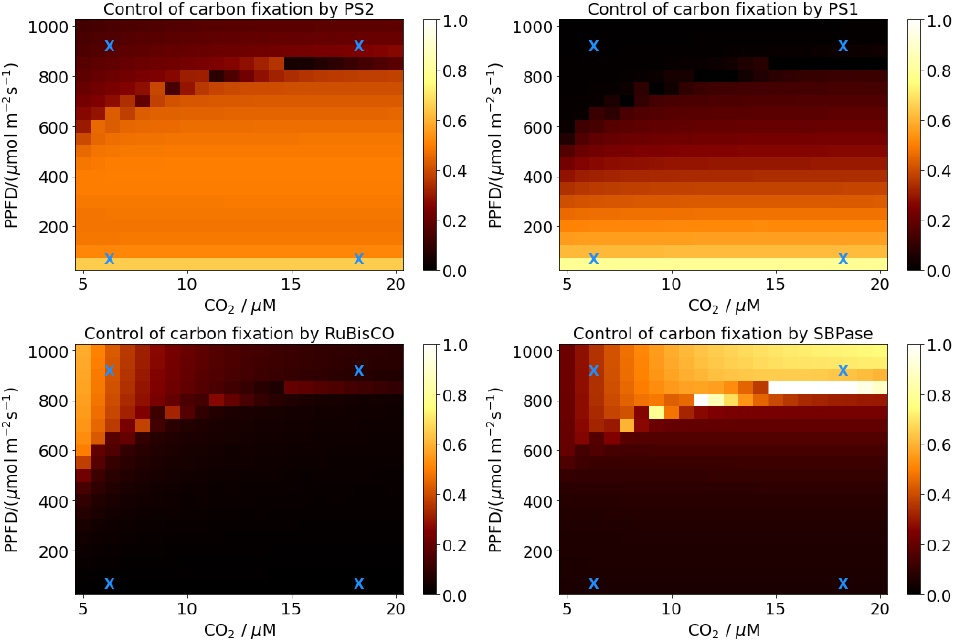
The flux control of PSII (top left), PSI (top right), RuBisCO (bottom left), and SBPase (bottom right) on carbon fixation in light intensities ranging from 50 to 1000 μmol m^−2^ s^−1^and in CO_2_ concentrations ranging from 5 to 20 μM. The control coefficients are indicated by the heat map with dark areas indicating low and light areas high control. The blue crosses are the reference points for the further analyses indicating low CO_2_ /low light (6 μM, 100 μmol m^−2^ s^−1^), low CO_2_ /high light (6 μM, 900 μmol m^−2^ s^−1^), high CO_2_ /low light (18 μM, 100 μmol m^−2^ s^−1^), and high CO_2_ /high light (18 μM, 900 μmol m^−2^ s^−1^).

In the low light regime, both photosystems have substantial control whereas the CBB cycle enzymes exert almost no control. This changes drastically for higher light intensities, where PSI exerts practically no control and PSII only a small but distinguishable control, whereas the CBB cycle enzymes now control the carbon fixation rate. In high light, a gradual shift in control from RuBisCO to SBPase can be observed as CO_2_ concentrations increase. SBPase has the highest control in high light intensities and high CO_2_ concentrations, while RuBisCO is the dominant reaction in high light intensities and low CO_2_ concentrations. High control of RuBisCO in high light and low CO_2_ concentrations has also been found experimentally [28].

Summarizing, this analysis shows that the control on carbon fixation switches from photosystem I in low light to photosystem II in medium light intensities to SBPase and RuBisCO in high light intensities, where RuBisCO control dominates in low and SBPase control in high CO_2_ concentrations. For our further analyses, we define four reference conditions for low CO_2_ / low light (6 μM, 100 μmol m^−2^ s^−1^), low CO_2_ / high light (6 μM, 900 μmol m^−2^ s^−1^), high CO_2_ / low light (18 μM, 100 μmol m^−2^ s^−1^), and high CO_2_ / high light (18 μM, 900 μmol m^−2^ s^−1^). These conditions are indicated by blue crosses in Fig. 2.

### Control of photosynthetic intermediates

Besides the carbon fixation rate, also the states of the intermediates in the photosynthetic electron transport chain and the CBB cycle are important determinants for the efficiency and status of the photosynthetic system at large. In particular, poised redox levels of the electron carriers are indicative of the efficient functioning of the PETC, the concentrations of ATP and NADPH are important as ubiquitous redox and energy equivalents, and the CBB cycle intermediates must be above a certain level to ensure the cycle runs efficiently [17]. Moreover, various mechanisms ensure that, in particular in high light, photodamage by reactive oxygen species (ROS) is minimized.

The electron carriers behave as expected (see figures S3 to S6). In general, upstream reactions have a positive control on their redox state, while downstream reactions exert a negative control. For example, the redox state of plastoquinone is strongly positively controlled by PSII, slightly positive by the cyclic electron flow (which feeds back electrons from ferredoxin to plastoquinone), and negatively or not at all by downstream processes, such as PSI or the CBB enzymes (see figure S3). The only electron carrier that is more reduced when the CBB cycle enzymes are increased is plastocyanin, which is in agreement with previous model analyses [26] (Fig. S4). An interesting observation is that under low light, both ferredoxin and NADPH are less reduced if PSI activity is enhanced, although ferredoxin is a direct product of PSI (Figs. S5 and S6). A possible explanation for this counter-intuitive finding is that the cyclic electron flow is strongly increased with increasing PSI activity (Fig. S7) and that, together with the increased CBB activity (see above) this leads to a slight reduction of these two electron carriers. The control of ATP levels is complex (Fig. S8). For example, increased PSII leads to reduced ATP levels in very low light, increased in intermediate light (still below the quencher saturation threshold), and a slight reduction again for high light conditions. However, steady state energy levels range between 0.6 and 0.8 (fraction of ATP in the adenosine phosphate pool), which are in the range of measured values [29]. An interesting effect is observed when calculating the control on the total phosphates in CBB intermediates. Apparently, enhancing the fixation process (RuBisCO) leads to a reduction in CBB intermediates, whereas enhancing the recycling phase (SBPase) leads to an increase, except for very low light intensities (Fig. S9). ROS (simulated as stationary H_2_O_2_ concentrations) levels respond as expected. Increasing the photosystems leads to higher levels, while increasing cyclic electron flow, the b_6_f complex activity or the CBB cycle lead to reduced levels (Fig. S10.)

### Robustness of the control on carbon fixation in multiple environmental conditions

Control coefficients quantify the strength of control of individual processes in a metabolic network. They are system-wide properties and, as such, depend on the specific values of the kinetic parameters of the involved enzymatic reactions. Therefore, they should not be considered a rigid value independent of all choices in the model-building process or of varying external conditions. The control on RuBisCO, an essential enzyme for carbon fixation, by other reaction steps in the CBB cycle, PETC, or ASC-GSH cycle, is interesting for broadening our understand-ing of sequestering carbon in photosynthetic organisms.

In order to determine if the previously observed shift in control (Fig. 2) is a consequence of the structural design of the PETC and the CBB, we performed a robustness analysis. For this, we varied parameters by multiplying a randomly selected factor between 1/2 and 2 to generate 10000 perturbed parameter sets. For each parameter set, we analyzed the control exerted by PSII, cytochrome b_6_f, RuBisCO, FBPase, and SBPase on carbon fixation in the four reference conditions for low/high CO_2_/light as defined above. Figure 3 shows the distributions of flux control coefficients on carbon fixation by selected reactions. With most parameter sets, the photosystems had a much higher control on carbon fixation in low light intensities in both CO_2_ concentration conditions than reactions in the CBB cycle. In low light intensity, cytochrome b_6_f and RuBisCO have almost no control, and SBPase, while detectable, has only a minor in-fluence. Investigating the correlation of control coefficients for both photosystems under low-light conditions reveals that these two processes indeed share the main flux control in a proportion, which depends on the exact parameter values (Fig. 4).

**Figure 3.**
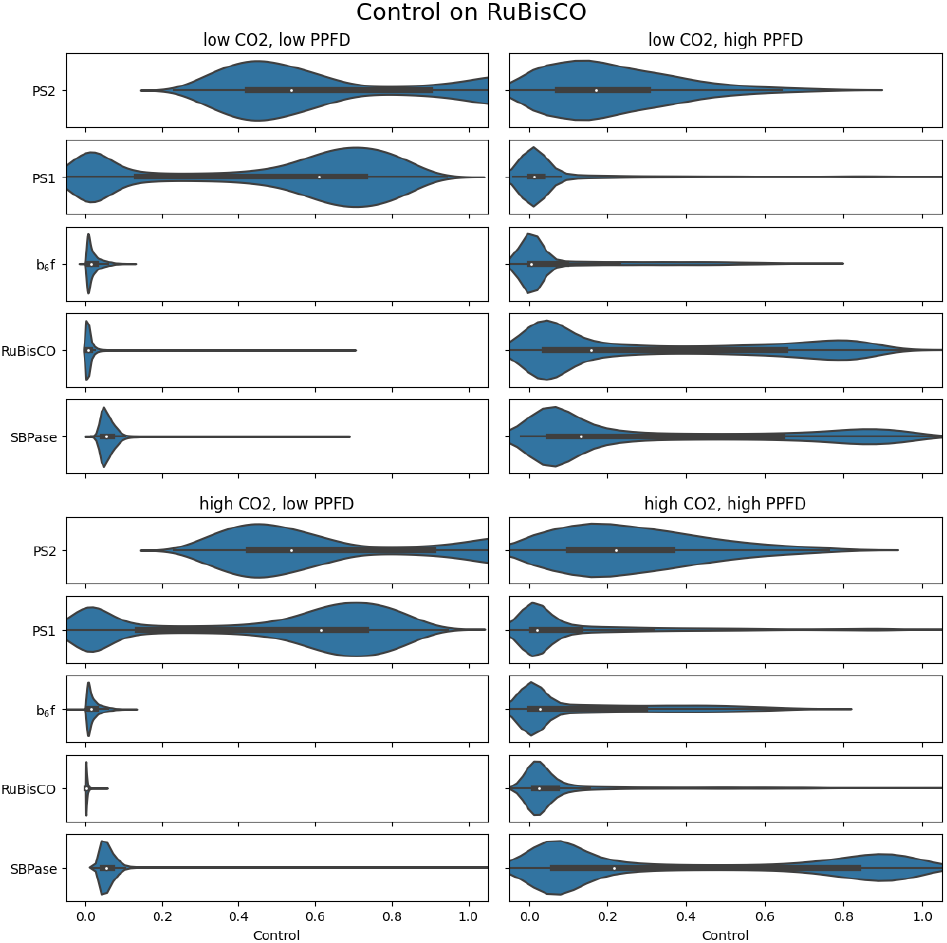
Distribution of flux control of key reactions in the PETC and CBB cycle on carbon fixation over 10000 sets of randomly perturbed parameters with a factor between 1/2 and 2.

**Figure 4.**
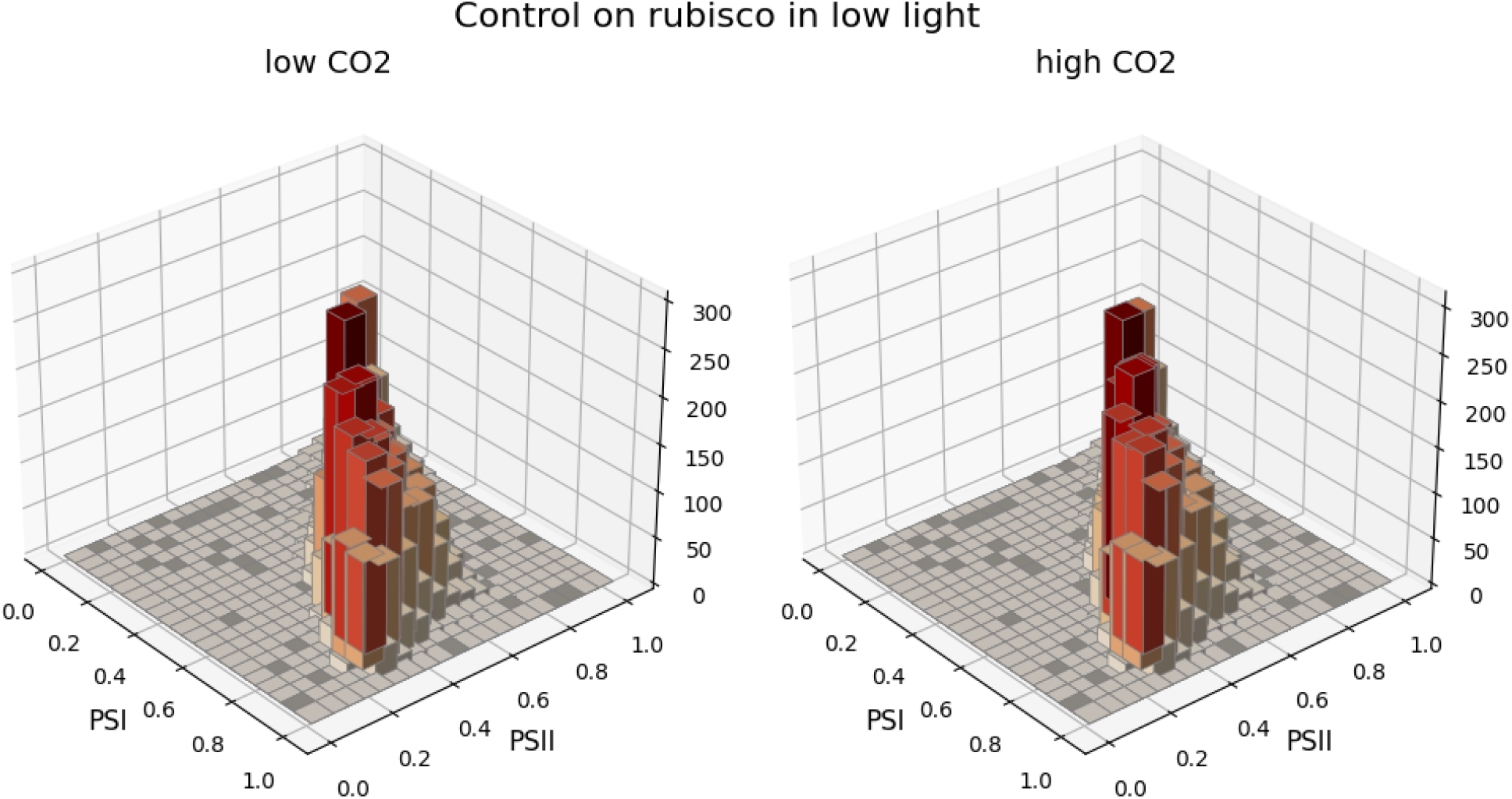
Correlation of control coefficients for both photosystems on carbon fixation under low-light conditions represented as 3D histogram. The z-axis indicates the amount of control coefficient in a specific numerical range. The calculation is based on 10000 randomly generated parameter sets as described in the text.

The control of photosystems on carbon fixation is drastically reduced in high light conditions. Especially photosystem I has lost almost all its control. As a general trend, the distribution of flux control coefficients in high light is broader than in low light. The main controlling steps are now on the demand side of photosynthesis, in particular on RuBisCO and SBPase. SBPase is, besides photosystem II, the controlling reaction for carbon fixation in high light intensity and high CO_2_ conditions. At the same time, RuBisCO is the main factor in high light intensity and low CO_2_ concentrations. Correlating the control coefficients of these two central CBB enzymes shows for most randomly selected parameter sets RuBisCO does not exert any considerable control under high light and high CO_2_ concentrations, while under high light but low CO_2_ the control can be on either of these enzymes (or none of the two) but only for a few parameter sets the control is shared between the two enzymes (Fig. 5).

Overall, our robustness analysis, in which we randomly varied parameters by up to a factor of two, confirms our previous observations, namely that the control shifts from the photosystems in low light to the CBB cycle enzymes in high light. Under the latter conditions, RuBisCO exerts a higher control if ambient CO_2_ concentrations are low, while SBPase is the controlling step under high CO_2_ concentrations. This indicates that the shift in control is less a kinetic, parameter-dependent, effect but rather a structural property of photosynthesis.

## Discussion & Conclusions

Photosynthesis is a supply-demand system. The supply (PETC) and demand (CBB cycle) sides must be coordinated to ensure efficient photosynthesis. Considering the often rapidly and unpredictably changing light intensities [16] plants are exposed to in natural environments, maintaining such a coordination appears challenging. It is plausible to assume that the present environmental-dependent regulatory mechanisms controlling carbon fixation have evolved to be highly efficient, considering the direct effect that carbon capture has on plant growth and fitness. This work presents an *in silico* analysis of the control over carbon fixation in different environmental conditions. For this, we used a published model of photosynthesis that combines the supply side (PETC), the demand side (CBB cycle), and the Ascorbate-Glutathion cycle [26].

Such a supply-demand photosynthesis model allows quantifying the control that individual processes have on the overall carbon fixation rate. We focused in particular on photosystems I and II, cytochrome b_6_f, RuBisCO, and SBPase, which were reported to exert control on carbon fixation under several conditions [21, 13, 23]. Using Metabolic Control Analysis, we quantified the control of these single steps on carbon fixation for different simulated environmental conditions. By simultaneously varying the light intensity and CO_2_ concentration, we could show that the control shifts from the photosystems in low light intensities to RuBisCO and SBPase in high light intensities but then from RuBisCO in low to SBPase in high CO_2_ concentrations (Fig. 2). The shift of the control confirms that most of the reactions previously reported to control the flux are indeed critical for regulating carbon fixation. However, whether PSII, PSI, RuBisCO, or SBPase is the main controlling factor strongly depends on the external conditions. In our photosynthesis model, a relatively sharp threshold marks the transition between a supplyand a demandcontrolled situation (see Fig. 2). This threshold, separating ‘low’ and ‘high’ light conditions, occurs when the quenching mechanism reaches its maximal capacity (see Fig. S2). This results in a reduction of most electron carriers and a sharp accumulation of protons in the lumen. The PETC still operates at a fast rate, so that ATP and NADPH production is no longer limiting for carbon fixation. It is an open question whether this sharp transition is a feature of the specific model that was used for this analysis or whether this is actually a systemic property of photosynthesis. We assume that the transition from non-saturated to saturated quencher is not as sharp *in vivo* as suggested by the model, but that the principle feature, namely that high light intensity results in a shift of control to the demand reactions of the CBB cycle, is a structural feature of the photosynthetic supply-demand system. The continuous transition under high light between the RuBisCO and SBPase-mediated control suggests that at high CO_2_ concentrations, the carboxylation by RuBisCO is not determining carbon fixation rate, but rather the distribution of the intermediates in the CBB cycle through SBPase. To test whether the shift in control is a kinetic property of the rates in photosynthesis or follows from the structure resulting from the interconnections of the PETC and CBB cycle, we performed a robustness analysis by randomly varying kinetic parameters by a factor between 0.5 and 2. Figure 3 illustrates that, at least in our model representation, the control shift is a property that occurs with many parameter sets. This observation indicates that the shift of control indeed seems to be a structural feature, and rather independent on the specific parameter values.

Interesting patterns emerge by correlating the flux control coefficients obtained by the robustness analysis. In the low light regime, the control is shared mostly among the two photosystems (Fig. 4), where often one of the photosystems exhibits a higher control than the other. Which photosystem exerts the higher control apparently depends on the specific numerical parameter values. Additionally, the fact that both photosystems always have clearly non-zero control for all parameter sets in low light underlines the importance of the PETC for carbon fixation as a limiting factor in these conditions. The light-driven photosystems ultimately determine the flow of electrons through the PETC and the translocation of protons into the lumen, hence the production of ATP and reduction equivalents required by the CBB cycle. Correlating the control coefficients quantifying the importance of RuBisCO and SBPase under high light shows a drastically different picture. Fig. 5 reveals that typically carbon fixation is either controlled by RuBisCO or by SBPase, but the control is rarely shared. This is especially pronounced in high light intensity and low CO_2_ concentration conditions. Fig. 5 also reveals that for a substantial number of parameter combinations, neither RuBisCO nor SBPase exerts control over the carbon fixation rate. A closer inspection reveals that in these cases, the control is, in fact, on the photosystems. In fact, correlating the total control (sum of control coefficients of the individual processes) of the supply reactions with the total control of the demand reactions reveals that the control lies either on the supply side or on the demand side, but is rarely shared between both sides (Fig. S12). An interesting observation is that even in high light conditions, the model exhibited control by the light reactions for a substantial fraction of parameter sets. A possible explanation for the observation that also in high light for many parameter sets the control lies on the photosystems is that variations of the parameters can lead to scenarios where our selected ‘high light’ condition is actually not perceived as saturating light. In order to test this hypothesis, we relate the control exhibited by the dark reaction to the simulated stationary lumen pH (Fig. S14). This analysis shows that whenever the control is on the dark reactions, the pH is low, indicating that light (and the quencher) is saturated, whereas low control by the dark reactions is associated with a high lumenal pH, indicating nonsaturating conditions.

**Figure 5.**
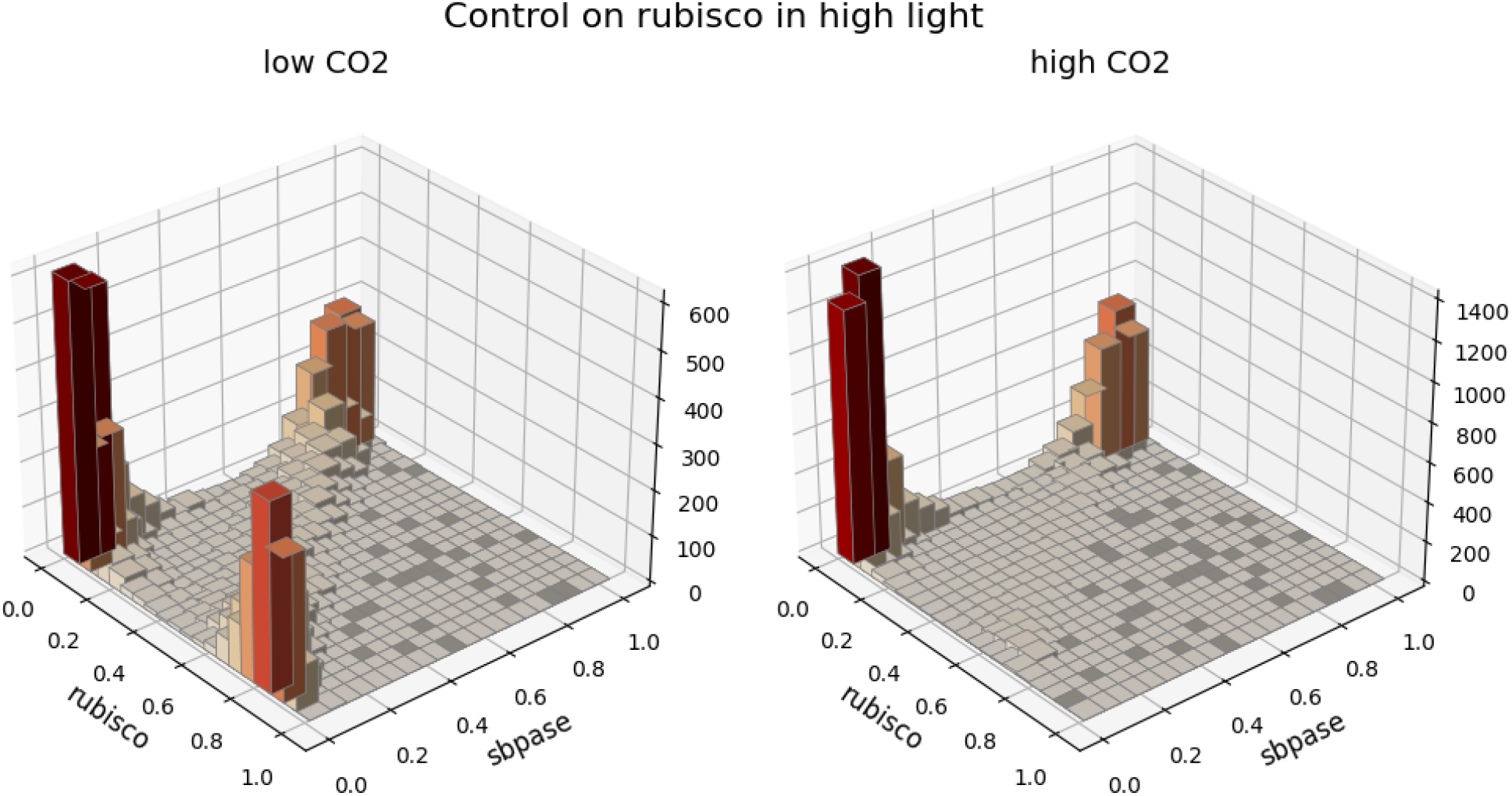
Control coefficients of carbon fixation by RuBisCO vs. SBPase under high-light conditions represented as 3D histogram. The z-axis indicates the amount of control coefficients in a specific numerical range. The calculation is based on 10000 randomly generated parameter sets as described in the text.

Some experimental and theoretical studies claim that cytochrome b_6_f controls the photosynthetic flux [27, 13]. In contrast, our analysis suggests that cytochrome b_6_f exhibits a considerable control only for very few parameter sets (Fig. 3). However, when we systematically decrease the activity of cytochrome b_6_f, also in our model cytochrome b_6_f can become a rate-controlling step (Fig. S11). These considerations show that seemingly conflicting reports on the control of cytochrome b_6_f are not necessarily contradictory. In fact, the parameters describing the composition and kinetic properties of the photosynthetic ap-paratus have an important influence on the strength of control.

Most concentration control coefficients behave as expected. For the electron carriers, upstream reactions exert positive and downstream reactions negative control. We obtained an initially counter-intuitive result only for ferredoxin and NADPH, as they are both less reduced when PSI activity is enhanced in low light. This observation might be explained through an increased cyclic electron flow with a concomitant increase in CBB cycle activity. The cyclic electron flow is an integral part of the photosynthetic machinery adjusting the ATP/NADPH ratio in the PETC and, hence, is an essential regulatory mechanism. Responding to the ADP/NADPH ratio required by the demand reactions, effects of other processes can be reduced or even reverted, when compared to a system without cyclic electron flow. The regulatory effects of CEF may also explain the complex patterns in the control that some processes have on ATP concentrations. These results demonstrate that control in a complex system is often non-trivial, and altering reaction rates may result in counter-intuitive effects.

Exploring a previously published supply-demand photosynthesis model with metabolic control analysis, we could resolve seemingly contradictory statements about which reactions have the strongest control on carbon fixation. We showed that basically all reactions previously reported to exert a strong control can indeed have high flux control under some conditions. It is important to note that all results have been obtained from a single, imperfect model. The model does not, for example, include the important process of photorespiration or stomatal aperture. It is unclear how far the interpretation of the results and the derived conclusions can be generalized. Still, the strength of theoretical analyses is that also with simplified and imperfect models, general features can be identified and novel hypotheses derived. For example, the general pattern observed in our analysis how the control shifts between key enzymes and complexes depending on light intensity and CO_2_ concentrations is plausible and generally applicable. By understanding the principles how regulation depends on environmental conditions, new data can be interpreted in a highly informed manner. With our study, we aimed at demonstrating the usefulness of systematic model analyses with Metabolic Control Analysis in understanding the metabolic regulation in complex networks.

## Methods

### Model description

For the *in silico* analyses, we used a previously published model of photosynthesis [26]. This model combines mechanistic descriptions of the ETC, and the CBB cycle, supplying and consuming ATP and NADPH. The model includes the regulation of CBB enzymes via thioredoxin and mechanisms responsible for producing and scavenging ROS around PSI. The scavenging of ROS is mediated by a module representing the ASC-GSH cycle.

### Metabolic control analysis

The flux 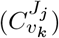 and concentration 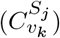 coefficients are defined, as

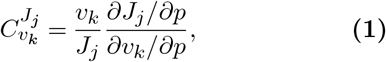

and

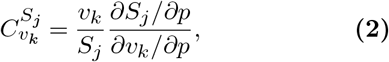

where the steady state fluxes and concentrations are denoted as *J*_*j*_ and *S*_*j*_, respectively. *p* is a kinetic parameter affecting only the reaction *k* with rate *v*_*k*_ directly. In the computational analyses the control coefficients were numerically approximated using central difference and varying the parameter *p* by *±*1%.

### Reaction dendrogram and Reaction correlation coefficients

A set of flux vectors satisfying the steady state condition,

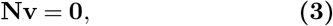

defines the nullspace of the stoichiometric matrix. A set of base vectors summarised in the kernel matrix **K**, in which they form the columns, span the nullspace. The kernel matrix can be obtained by the relation,

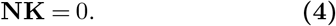

The reaction correlation coefficients were calculate following [22]. The kernel matrix was orthonormalized using the Gram–Schmidt process implemented in the sympy [18] package. For a pair of reactions and corresponding row vectors in the kernel matrix **k**_*i*_, and **k**_*j*_, the reaction correlation coefficients calculate as,

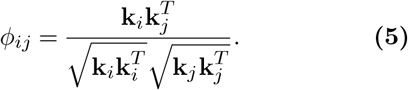

The dissimilarity matrix Δ_*ij*_, describing the angle between the row vectors of the kernel matrix **K**, was obtained using the reaction correlation matrix

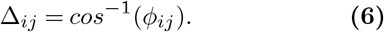

Hierarchical clustering, using Δ_*ij*_, was conducted with the WPGMA algorithm implemented in the scipy package [31].

### Robustness analysis

To analyze whether our previous results were due to general properties of the system or due to choice of parameters we performed a control analysis scan over 10000 sets of randomly perturbed parameters. The model parameters were randomly multiplied by factors between 1/2 and 2. This analysis was performed for low / high CO2 and light conditions.

## Abbreviations

MCA: Metabolic control analysis,
CBB: CalvinBenson-Bassham-cycle,
PETC: photosynthetic electron transport chain,
ROS: reactive oxygen species,
ASC-GSH: ascorbate-glutathione cycle,
SBPase: sedoheptulose-1,7-bisphosphatase,
RuBisCO: ribulose-1,5-bisphosphate carboxylase/oxygenase

## Author contributions

OE: initial idea and conceptualisation. OE: funding acquisition. MvA, TN, JE: visualisation. MvA, TN, JE: formal analyses. OE, TN, MvA: writing—original draft and introduction. TN, MvA: writing—original draft and methods. TN, MvA: writing—original draft and results. TN, OE: writing—original draft, discussion, and TN, MvA, OE, JE, NS writing—review and editing. All authors read and accepted the final version of the manuscript.

## Data Availability Statement

The original contributions presented in the study are included in the article/Supplementary Material, further inquiries can be directed to the corresponding author/s. The code can be found in under https://gitlab.com/qtb-hhu/photosynthesis-taskforce/2023-50-years-of-mca.

