## Supplemental Text for "What controls carbon sequestration in plants under which conditions?"

Table S1: Enzyme subsets

| Reaction abbreviations | Reaction names |
|---|---|
| 1. vG6P_isomerase, vPhosphoglucomutase, vStarch | glucose-6-phosphate isomerase, phosphoglucomutase, starch synthesis |
| 2. vdhap | dihydroxyacetone phosphate exporter |
| 3. vpga | 3-phosphoglycerate exporter |
| 4. vNDH | NADH reductase |
| 5. vMDAreduct | NADH:monodehydroascorbate oxidoreductase |
| 6. vgap | glyceraldehyde-3-phosphate exporter |
| 7. vE_activation, vE_inactivation, vFdTrReductase | ATP synthse activation and deactivation, Ferredoxin-Thioredoxin reductase |
| 8. vCyc | cyclic electron flow |
| 9. vLeak | proton leak over thylakoid membrane |
| 10. vSt12, vSt21 | state transition reactions |
| 11. vDeepox, vEpox | xanthophyll cycle reactions |
| 12. v10, v11, v12, v13, v8, v9, vF6P_Transketolase, vRuBisCO | transketolase (S7P reaction), ribose-5-phosphate isomerase, xylose-5-phosphate epimerase, ribulose-5-phosphate kinase, aldolase (SBP reaction), sedoheptulose bisphosphatase, transketolase (F6P reaction) , ribulose-1,5-bisphosphate carboxylase/oxygenase |
| 13. vAldolase, vFBPase | aldolase, fructose-1,6-bisphosphotase |
| 14. v3ASC, vDHAR, vGR | ascorbate:hydrogen peroxide oxidoreductase, glutathione:dehydroascorbate oxidoreductase, glutathione:NADP+ oxidoreductase |
| 15. vATPsynthase | ATP synthase |
| 16. vFNR | ferredoxin-NADP reductase |
| 17. vAscorbate, vMehler | |
| 18. vBPGA_dehydrogenase, vPGA_kinase | 1,3-bisphosphoglycerate dehydrogenase, phosphoglycerate kinase (PGA kinase) |
| 19. vPS2 | photosystem II |
| 20. vTPI | triose phosphate isomerase |
| 21. vB6f, vPS1 | cytochrome $b_6f$, photosystem I |
| 22. vLhcdeprotonation, vLhcprotonation | PsbS de- and protonation |
| 23. vEX_ATP | ATP export |
| 24. vEX_NADPH | NADPH export |
| 25. vFdred | |
| 26. vPTOX | PTOX |

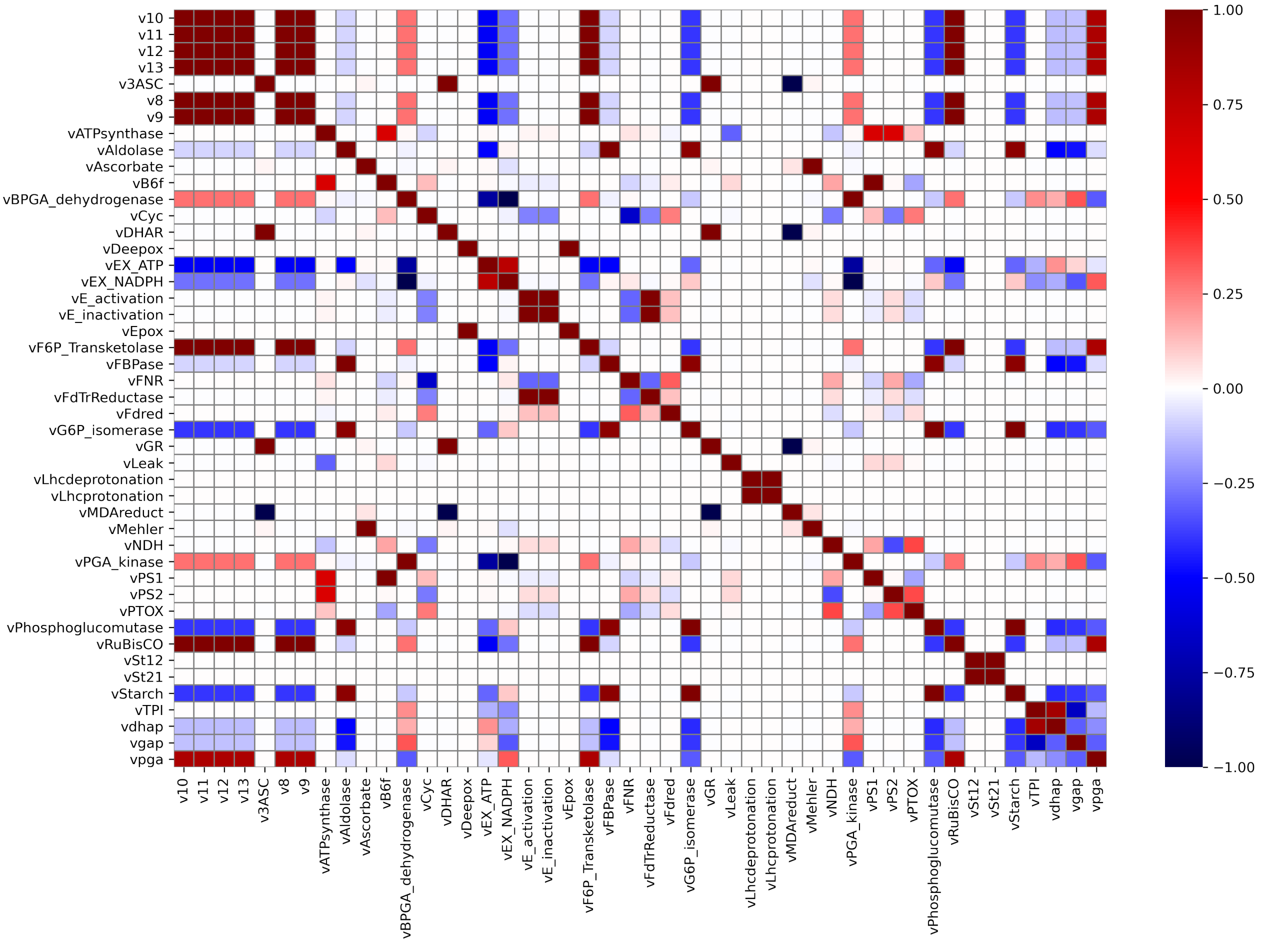


Figure S1: Reaction correlation coefficient matrix.

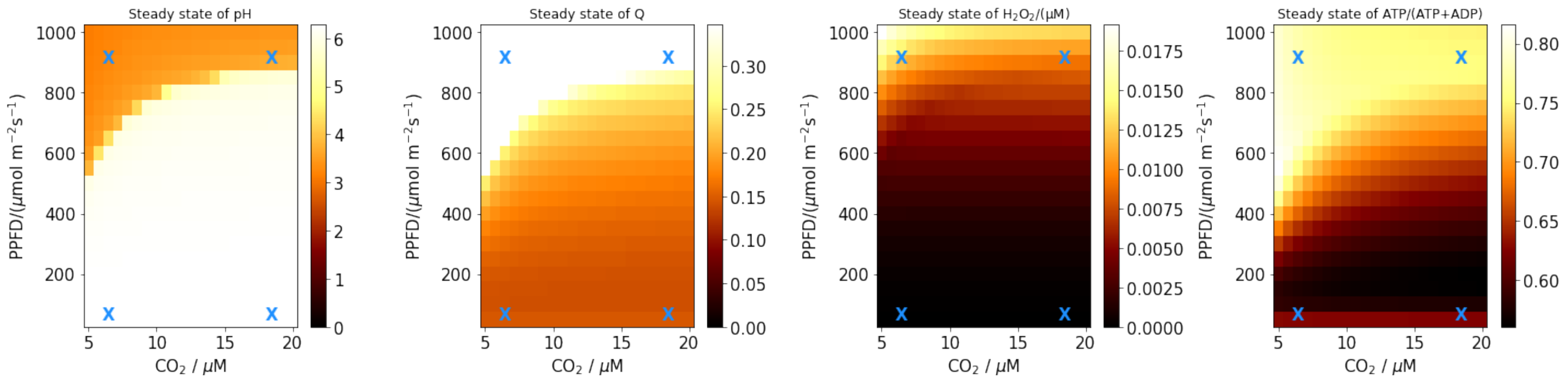


Figure S2: Steady state pH, quencher, $H_2O_2$, and ATP/(ATP+ADP) fraction.

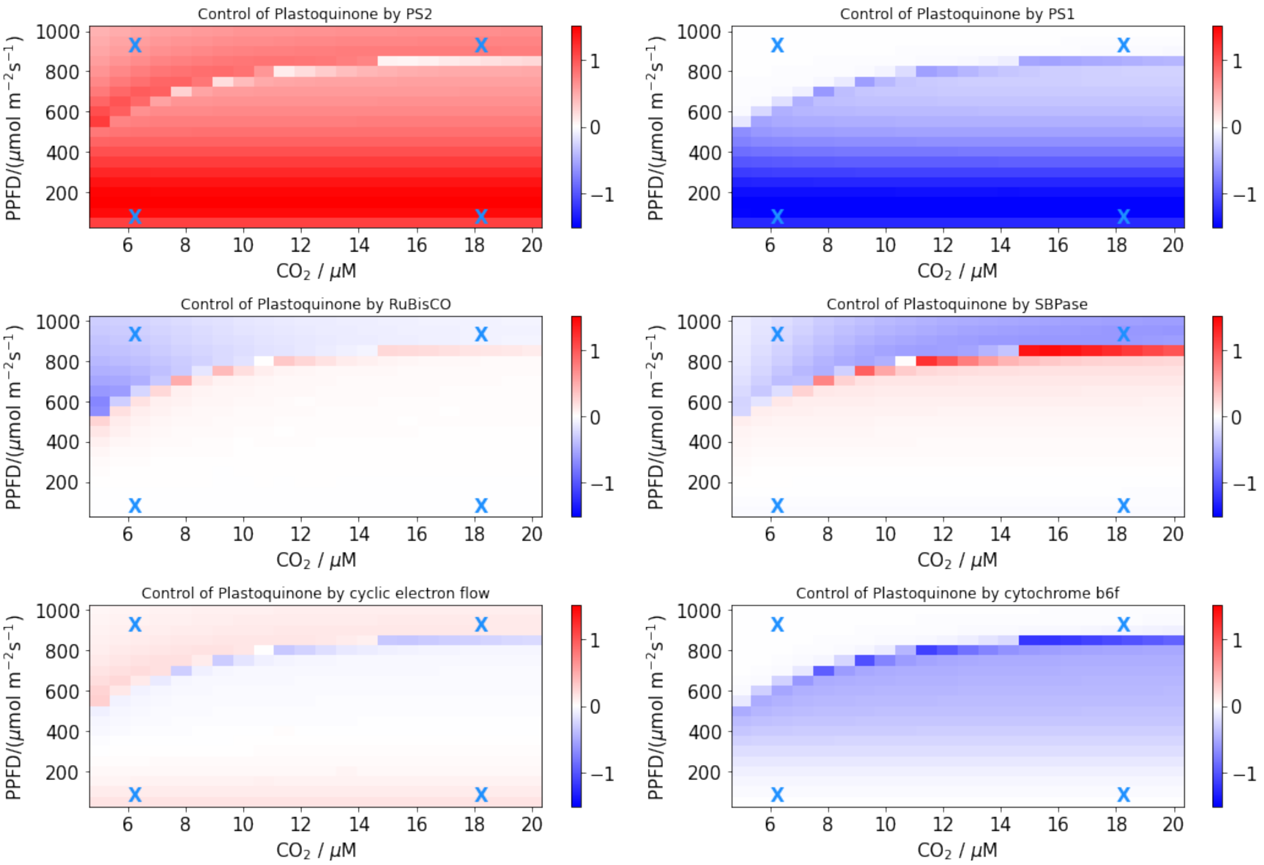


Figure S3: Concentration control coefficients of plastoquinone steady state concentration by photosystem II and I, RuBisCO, SBPase, cyclic electron flow, and cytochrome $b_6f$ under varying light intensities and carbon dioxide concentrations.

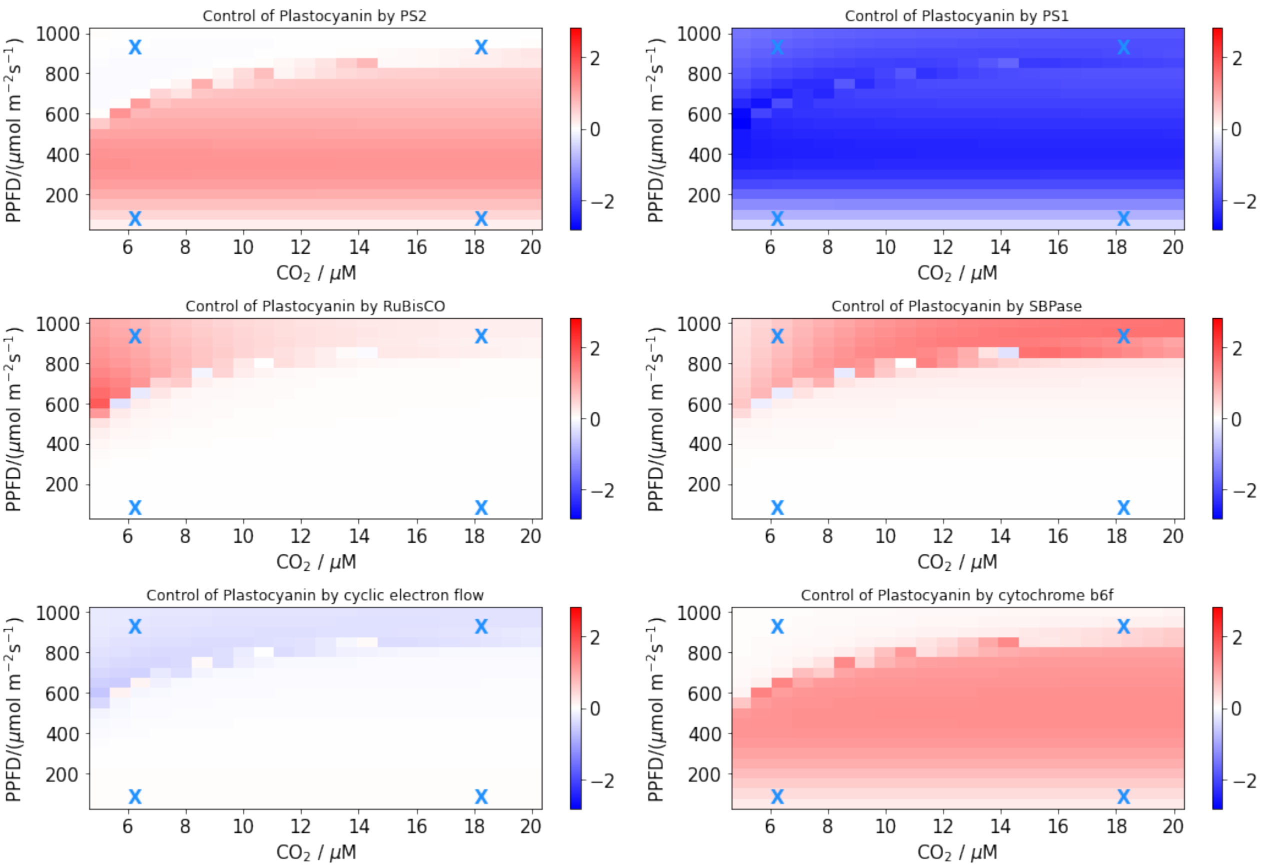


Figure S4: Concentration control coefficients of plastocyanin steady state concentration by photosystem II and I, RuBisCO, SBPase, cyclic electron flow, and cytochrome $b_6f$ under varying light intensities and carbon dioxide concentrations.

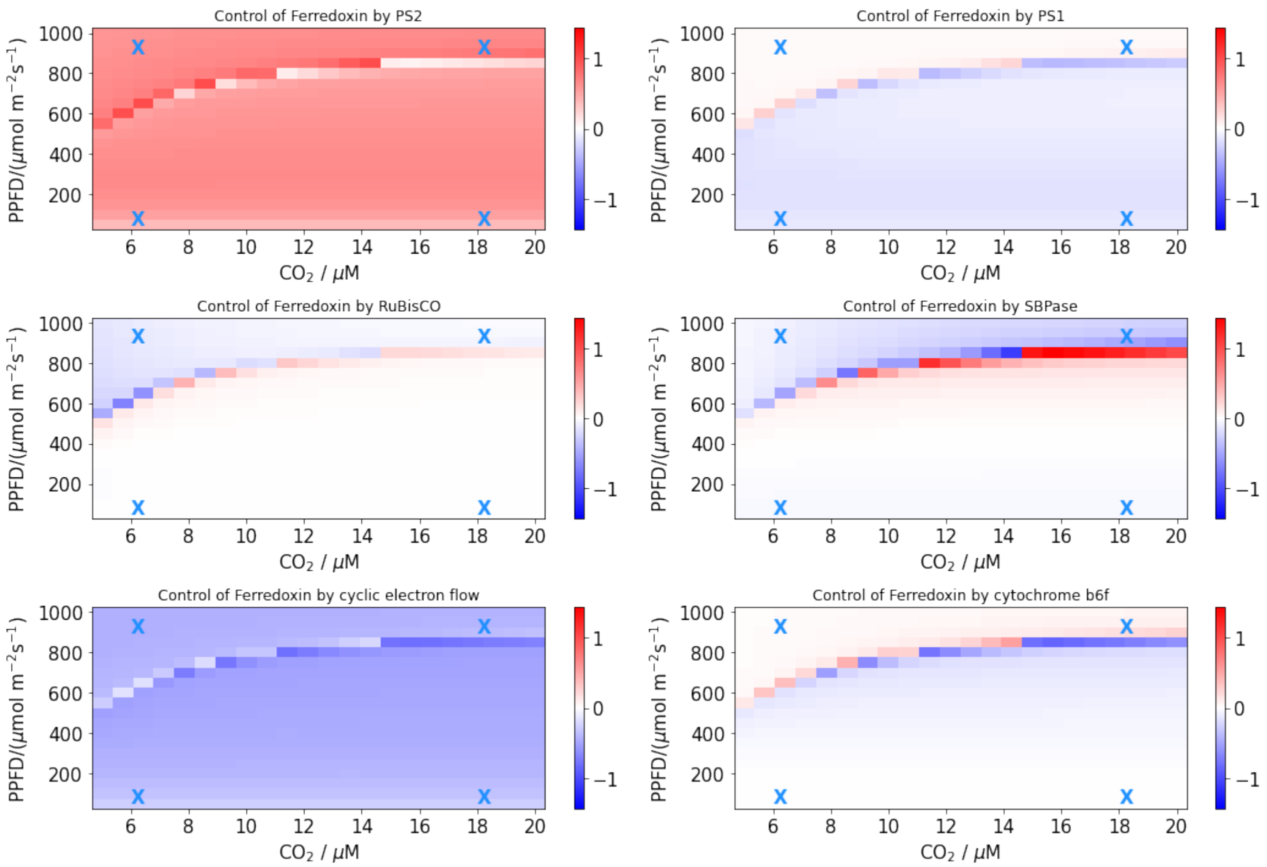


Figure S5: Concentration control coefficients of ferredoxin steady state concentration by photosystem II and I, RuBisCO, SBPase, cyclic electron flow, and cytochrome $b_6f$ under varying light intensities and carbon dioxide concentrations.

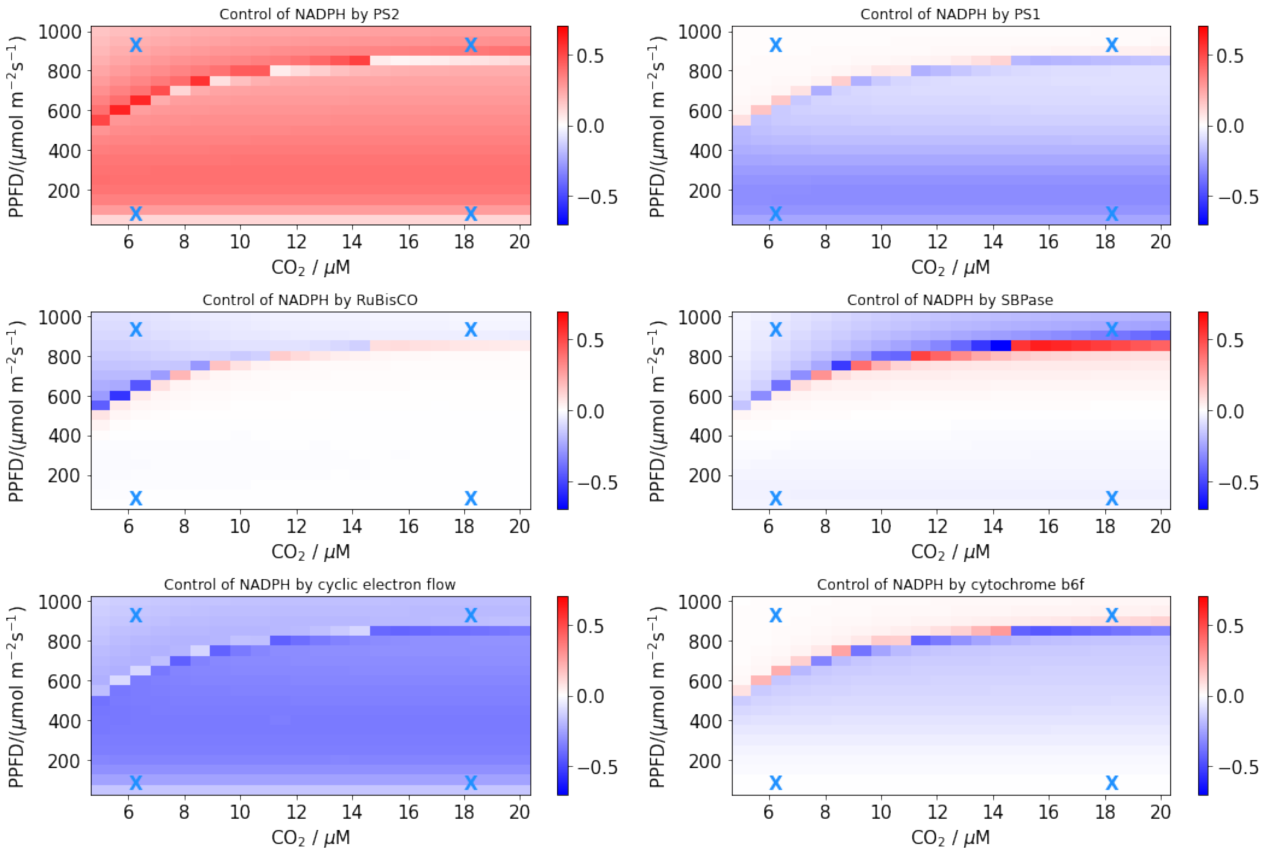


Figure S6: Concentration control coefficients of NADPH steady state concentration by photosystem II and I, RuBisCO, SBPase, cyclic electron flow, and cytochrome $b_6f$ under varying light intensities and carbon dioxide concentrations.

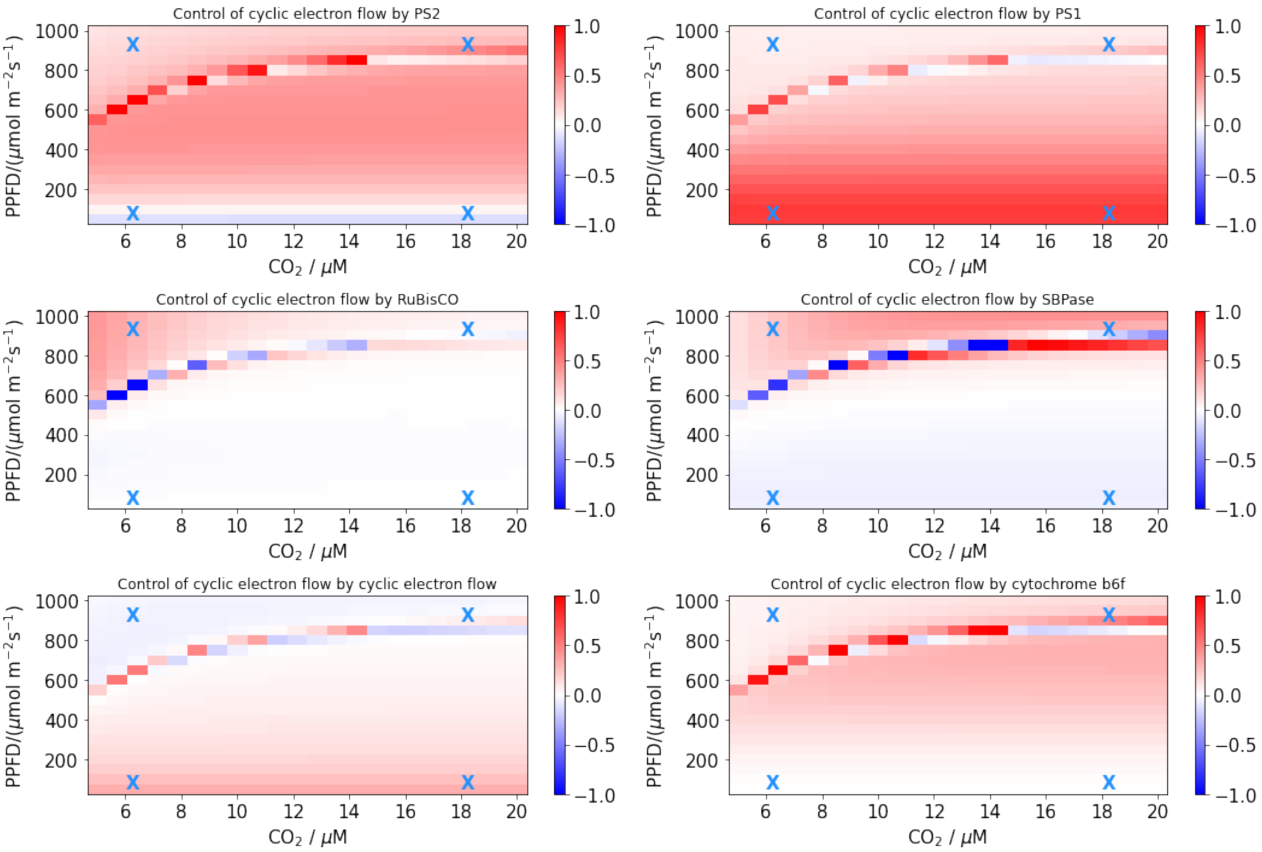


Figure S7: Flux control coefficients of cyclic electron flow by photosystem II and I, RuBisCO, SBPase, cyclic electron flow, and cytochrome $b_6f$ under varying light intensities and carbon dioxide concentrations.

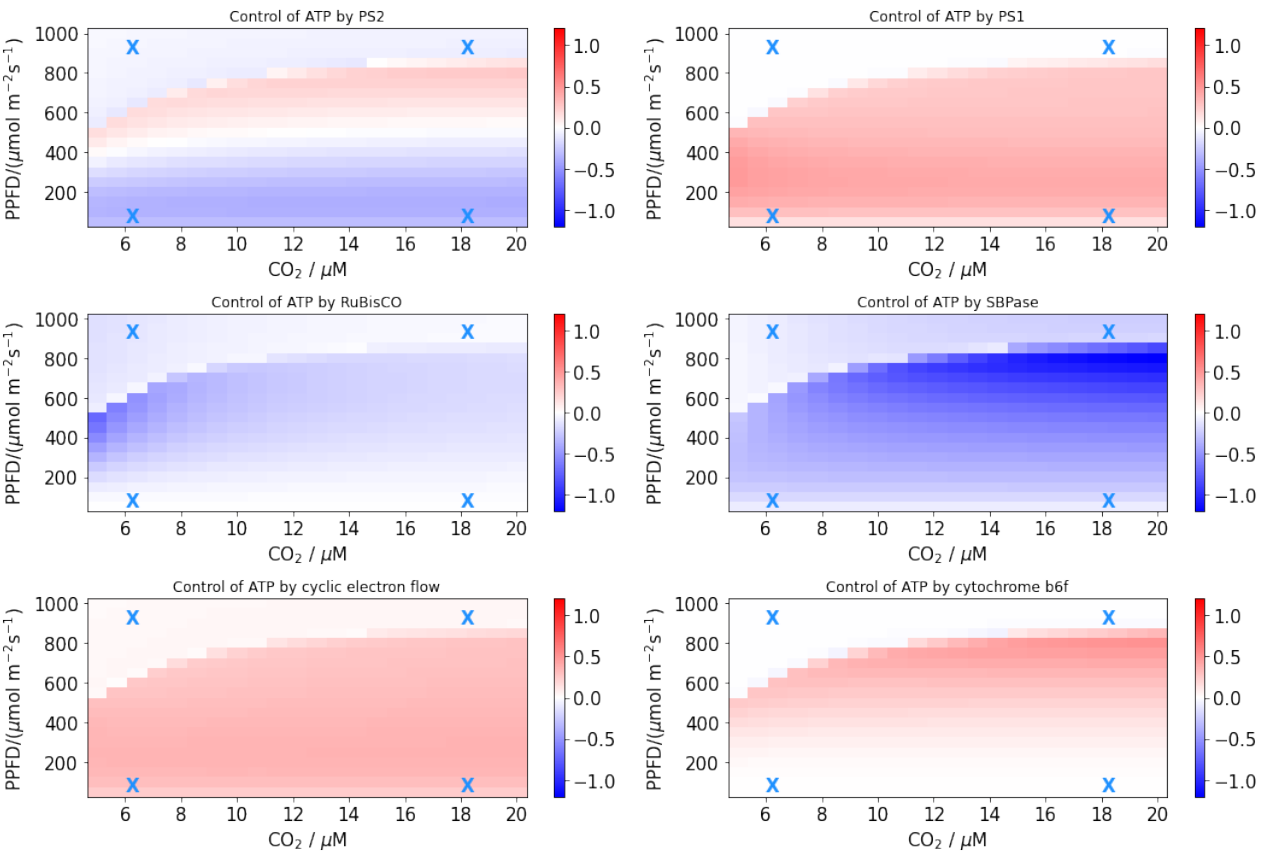


Figure S8: Concentration control coefficients of ATP by photosystem II and I, RuBisCO, SBPase, cyclic electron flow, and cytochrome $b_6$f under varying light intensities and carbon dioxide concentrations.

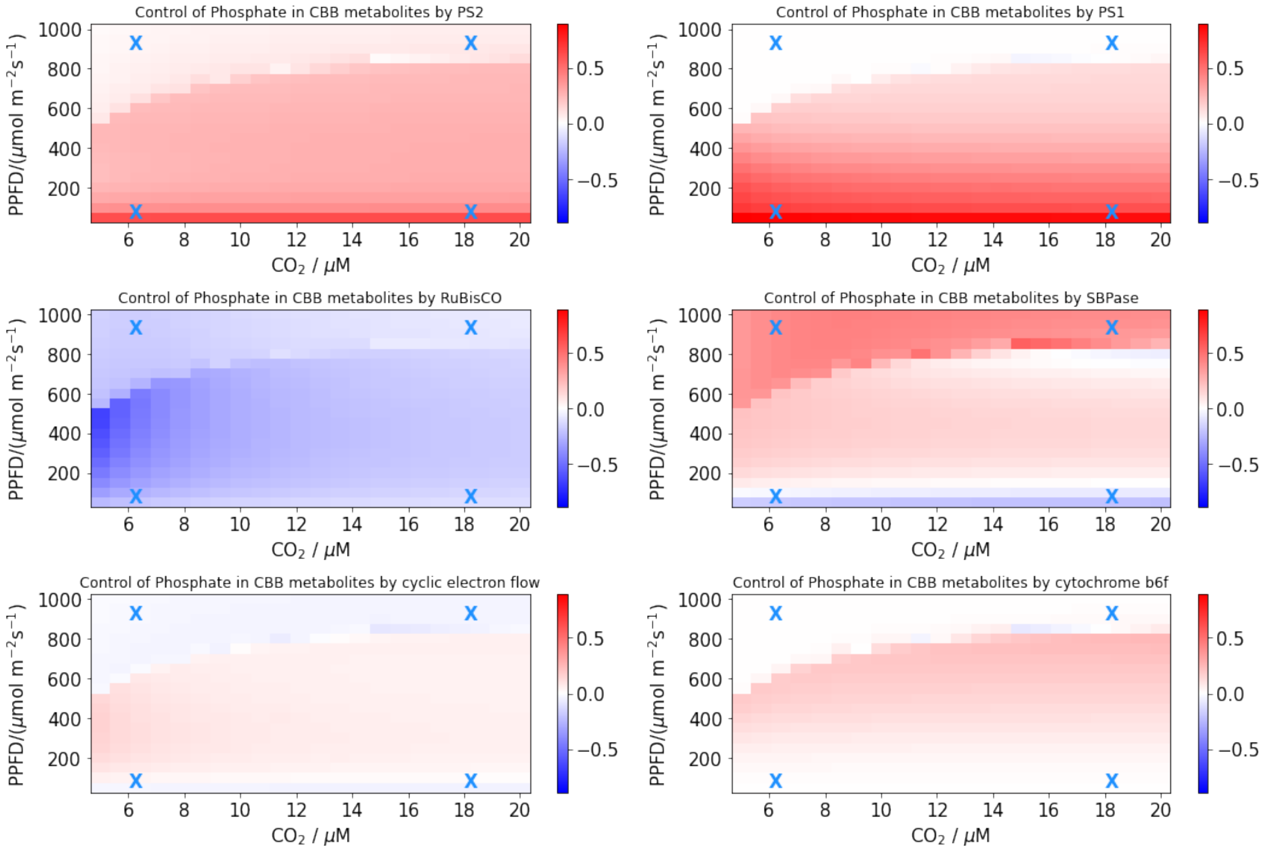


Figure S9: Concentration control coefficients of phosphate in CBB cycle intermediates by photosystem II and I, RuBisCO, SBPase, cyclic electron flow, and cytochrome $b_6f$ under varying light intensities and carbon dioxide concentrations.

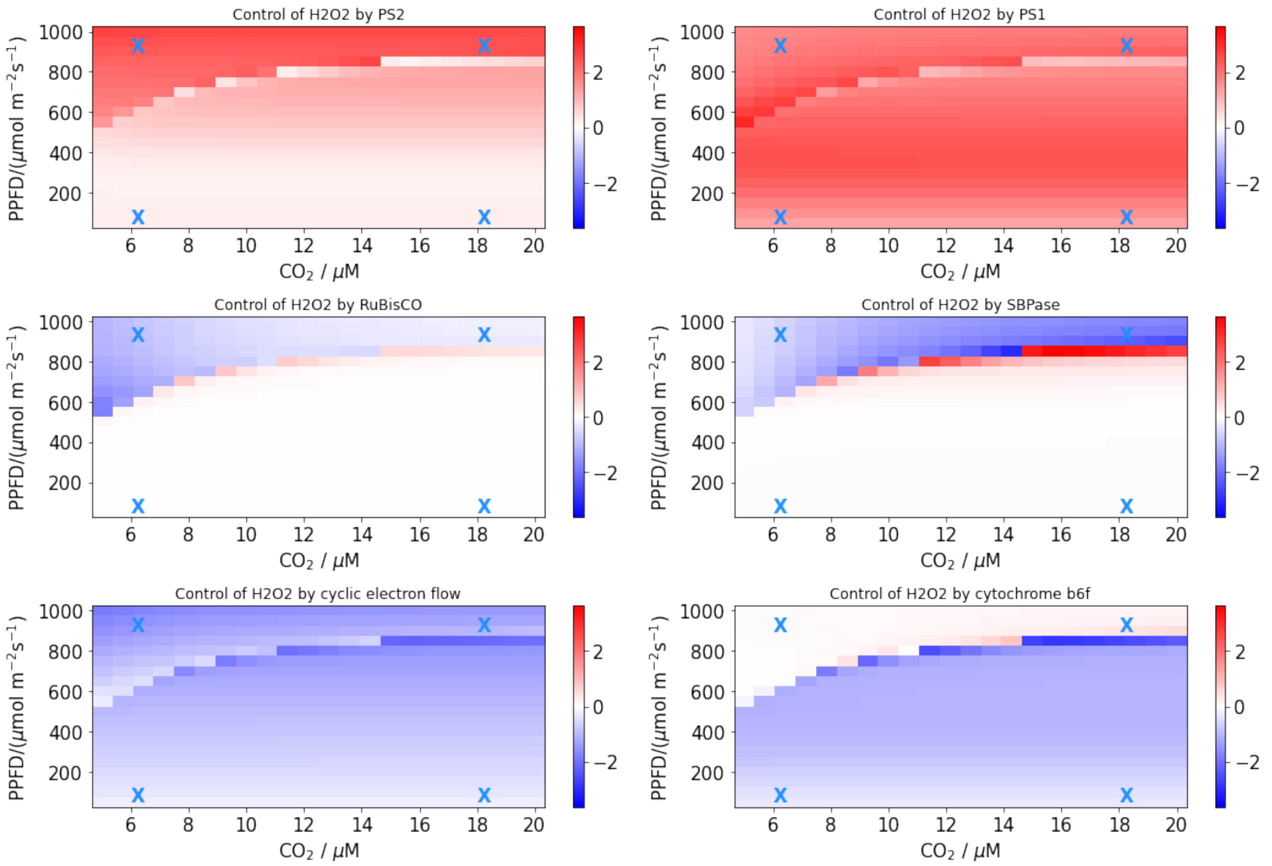


Figure S10: Concentration control coefficients of $H_2O_2$ steady state concentration by photosystem II and I, RuBisCO, SBPase, cyclic electron flow, and cytochrome $b_6$f under varying light intensities and carbon dioxide concentrations.

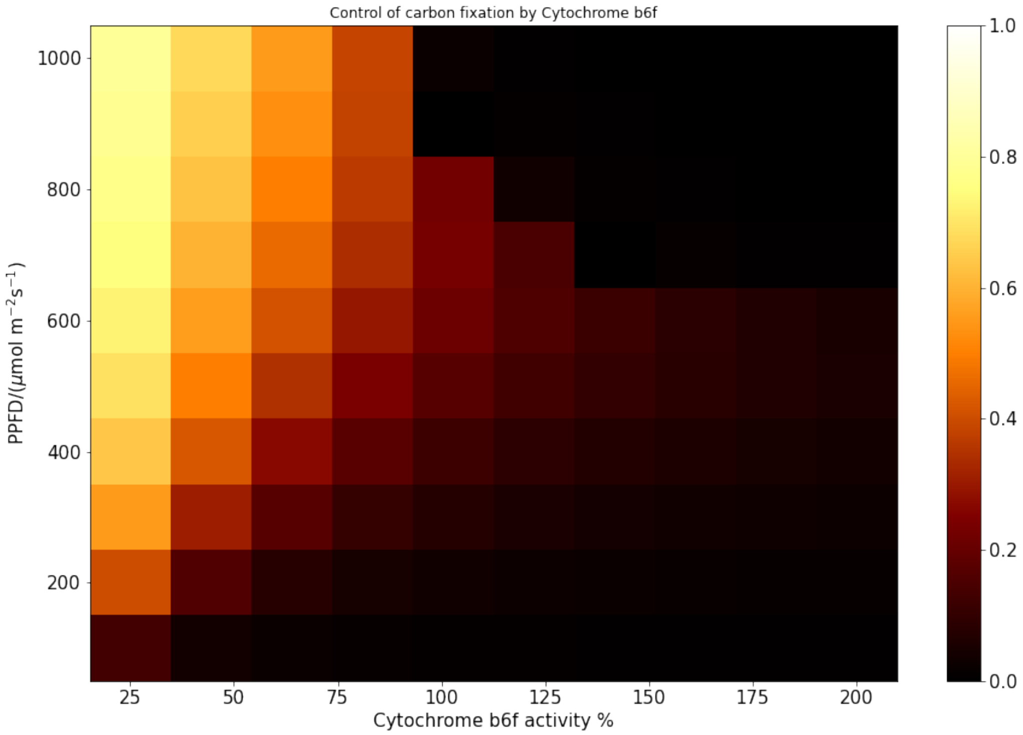


Figure S11: Flux control coefficients of carbon fixation by cytochrome $b_6$f under varying light intensities and cytochrome $b_6$f activities with high $CO_2$ conditions.

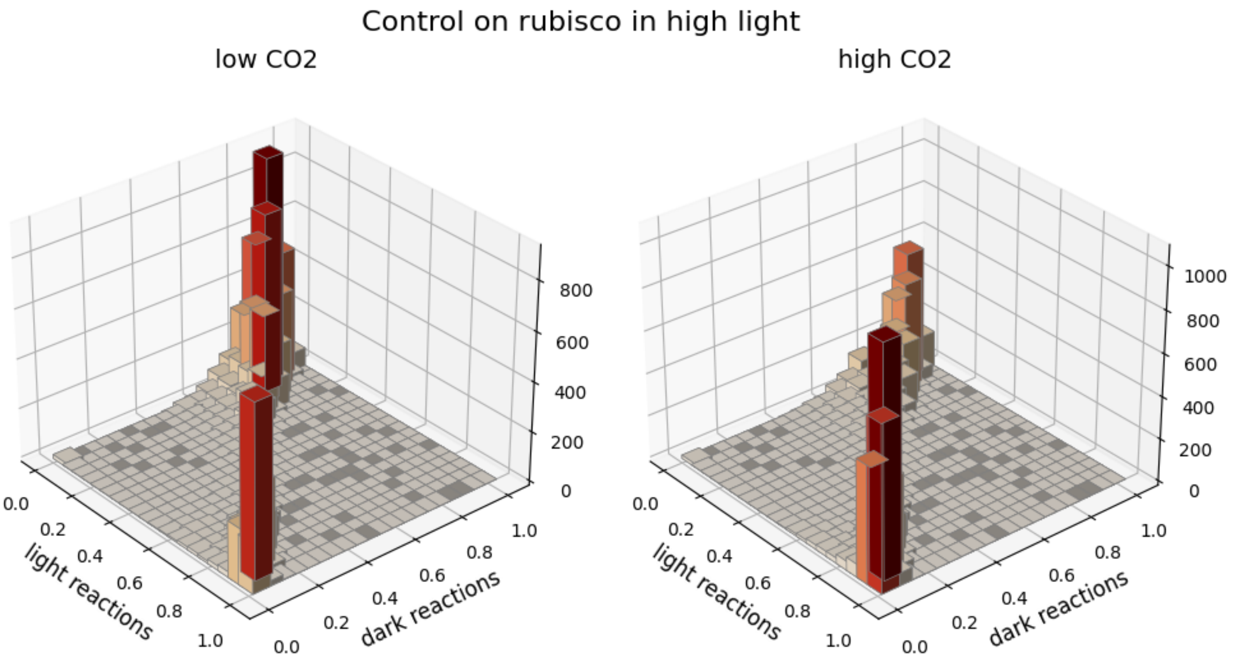


Figure S12: Control coefficients of carbon fixation by light vs. dark reactions in photosynthesis under high-light conditions represented as 3D histogram. The z-axis indicates the amount of control coefficients in a specific numerical range.

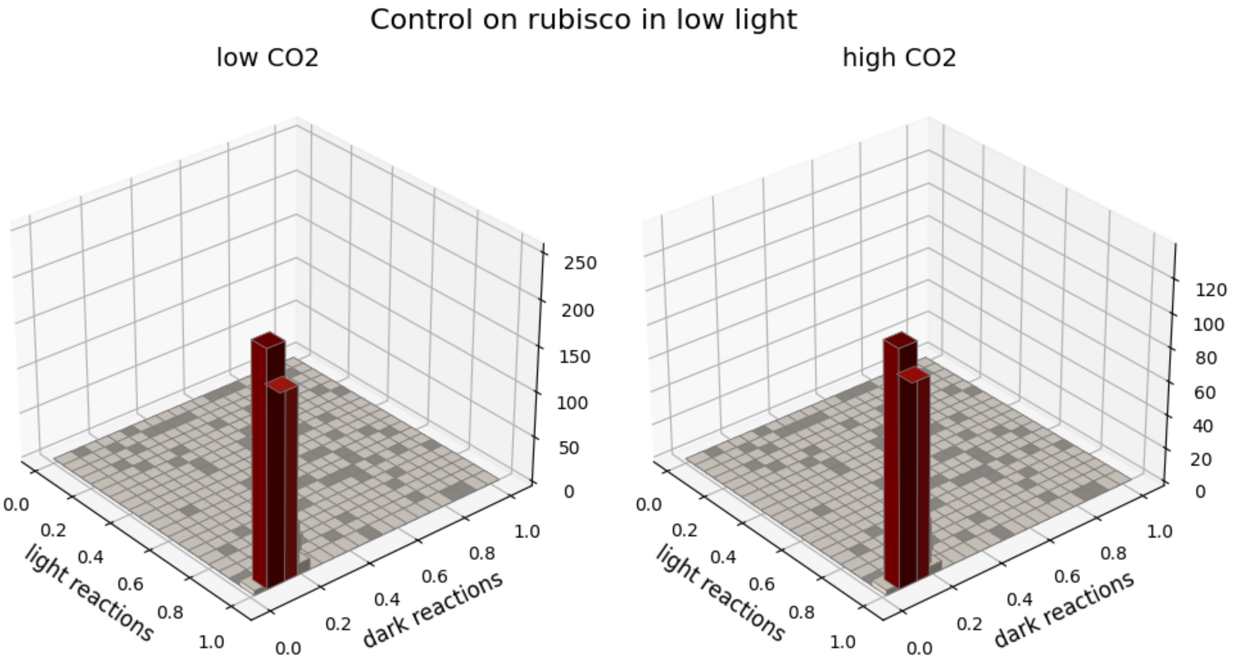


Figure S13: Control coefficients of carbon fixation by light vs. dark reactions in photosynthesis under low-light conditions represented as 3D histogram. The z-axis indicates the amount of control coefficients in a specific numerical range.

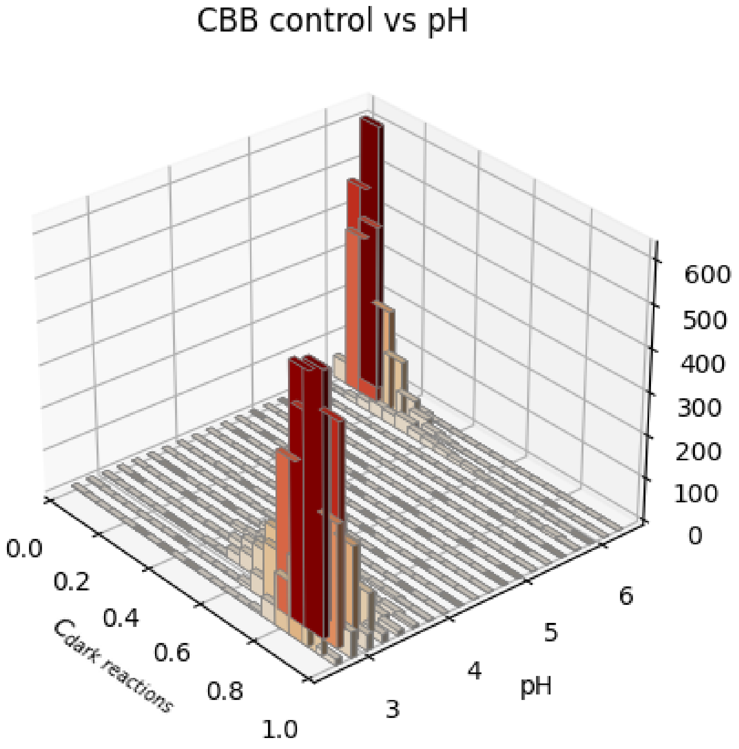


Figure S14: 3D histogram of steady state lumenal pH vs. control coefficients of dark reaction on carbon fixation. The z-axis indicates the amount of points in a specific numerical range.
